# A fast and sensitive size-exclusion chromatography method for plasma extracellular vesicle proteomic analysis

**DOI:** 10.1101/2024.01.17.576135

**Authors:** Ivo Díaz Ludovico, Samantha M. Powell, Gina Many, Lisa Bramer, Soumyadeep Sarkar, Kelly Stratton, Tao Liu, Tujin Shi, Wei-Jun Qian, Kristin E Burnum-Johnson, John T. Melchior, Ernesto S. Nakayasu

## Abstract

Extracellular vesicles (EVs) carry diverse biomolecules derived from their parental cells, making their components excellent biomarker candidates. However, purifying EVs is a major hurdle in biomarker discovery since current methods require large amounts of samples, are time-consuming and typically have poor reproducibility. Here we describe a simple, fast, and sensitive EV fractionation method using size exclusion chromatography (SEC) on a fast protein liquid chromatography (FPLC) system. Our method uses a Superose 6 Increase 5/150, which has a bed volume of 2.9 mL. The FPLC system and small column size enable reproducible separation of only 50 µL of human plasma in 15 minutes. To demonstrate the utility of our method, we used longitudinal samples from a group of individuals that underwent intense exercise. A total of 838 proteins were identified, of which, 261 were previously characterized as EV proteins, including classical markers, such as cluster of differentiation (CD)9 and CD81. Quantitative analysis showed low technical variability with correlation coefficients greater than 0.9 between replicates. The analysis captured differences in relevant EV-proteins involved in response to physical activity. Our method enables fast and sensitive fractionation of plasma EVs with low variability, which will facilitate biomarker studies in large clinical cohorts.

## Main text

Extracellular vesicles (EVs) are membrane-bound particles released by cells and play a role in tissue crosstalk [1]. EVs are classified based on their biogenesis and size. Exosomes are EV derived from multivesicular bodies ranging from 20 to 300 nm. Ectosomes, however, encompass a wide variety of EVs formed by directing budding from the plasma membrane, such as microvesicles (500-1000 nm) and apoptotic bodies (50-5000 nm) [1, 2]. EVs are composed of metabolites, nucleic acids, soluble and membrane proteins, as well as diverse types of lipids and saccharides [3]. EVs carry molecular signatures from their parental cell, presenting immense potential for biomarker development [3]. However, the purification step is a major hinderance to perform large-scale biomarker studies targeting EVs.

Current EV purification methods require large sample amounts, which is an impairment for most clinical studies. In addition, these methods have low performance, requiring methodological improvements. First, EVs share similar physicochemical properties with other biofluid components such as albumin and lipoproteins making EV purification difficult as they are minor components of biofluids, displaying one order of magnitude lower abundance than plasma lipoproteins [4]. Sequential purification steps can increase preparation purity, but result in high sample loss (≥99% after two purification steps) and are very time-consuming [5]. To better understand this issue, we recently performed a proteomics meta-analysis of EV datasets obtained with different purification methods, leading to the identification of 1717 proteins with high probability to be EV proteins [6]. With a better characterization of the plasma EV proteins and being able to distinguish them from contaminants, the purity of the samples became less important for quantitative proteomics analysis, and we focused on the sample recovery. By analyzing different methods, we found that size-exclusion chromatography (SEC) results in the highest EV recovery, averaging 35% of the proteome abundance [6].

We hypothesized that scaling down the required sample amount could result in similar EV-enrichment ratios as large-scale methods. We perform the analysis on a fast protein liquid chromatography system (ÄKTApurifier™, Cytiva) to improve the speed and reproducibility. To determine feasibility of this down-scaled approach, we injected 50 µL of commercially available human plasma (20-40 times less than most other methods) onto a Superose 6 GE increase 5/150 column (Cytiva) (**Figure 1A**). The elution was performed with an isocratic gradient of 10 mM Tris-HCl, pH 7.5 with 150 mM NaCl, 1 mM ethylenediaminetetraacetic acid, 0.02% sodium azide at 0.3 mL/min, resulting in remarkably similar elution profiles (**Figure 1B**). Samples collected into 10 fractions of 0.3 mL each, and incubated in the dark for 30 min with 1% sodium dodecylsulfate, 2 mM tris(2-carboxyethyl)phosphine and 1 mM iodoacetamide. Each sample was then precipitated with 80% acetone overnight at -20 °C and recovered by centrifugation at 4°C at 20.000xg for 30 minutes. Pellets were washed with cold acetone (-20 °C) and recovered by centrifuging for 10 minutes at 20.000xg and 4°C. They were dried at room temperature for 30 min, dissolved in 10% trifluoroethanol in 100 mM NH_4_HCO_3_, and digested with 1:100 (enzyme/protein ratio) endoproteinase LysC (Promega) for 2 hours at 37 °C with 800 rpm shaking and followed by 1:50 (enzyme/protein ratio) trypsin (Promega) for 16 hours at 37 °C with 800 rpm shaking. Peptides were analyzed in a Waters NanoAquity UPLC system (Waters Corporation) with a custom packed C18 column (4-cm x 150 μm inner diameter Phenomenex Jupiter) coupled with a Q-Exactive mass spectrometer (Thermo Scientific). Peptide separation was carried out with a gradient of water (solvent A) and acetonitrile (solvent B) both containing 0.1% formic acid (1-8% B in 2 min, 8-12% B in 18 min, 12-30% B in 55 min, 30-45% B in 22 min, 45-95% B in 3 min, hold for 5 min in 95% B and 99-1% B in 10 min). Eluting peptides were directly analyzed by nanoelectrospray ionization and the top 12 most intense parent ions were submitted to high-energy collision-induced dissociation fragmentation (2.0 m/z isolation width; 30% normalized collision energy; 17,500 resolution at 400 m/z), before being dynamically excluded for 30 s. Subsequent data processing with MSFragger in FragPipe v19.0 [7] against the human reference proteome (Uniprot Knowledgebase, downloaded January 2023), considered fully tryptic peptides, cysteine carbamidomethylation (fixed modification), and methionine oxidation and protein N-terminal acetylation (variable modifications). The analysis led to the identification 539 proteins with correlation between replicates ≥0.94 (**Figure 1C**). When compared to the EV proteins from our previous meta-analysis study [6], we detected a total of 95 EV proteins across the proteome (**Supplementary table 1**), and that they were significantly enriched with EV proteins (**Supplementary table 2**). We also assessed the distribution of EV across the fractions based on the classical markers, cluster of differentiation (CD)9 and CD8. These markers were only observed in fraction 3 (**Figure 1D**), suggesting that EV eluted exclusively in that fraction.

**Figure 1.**
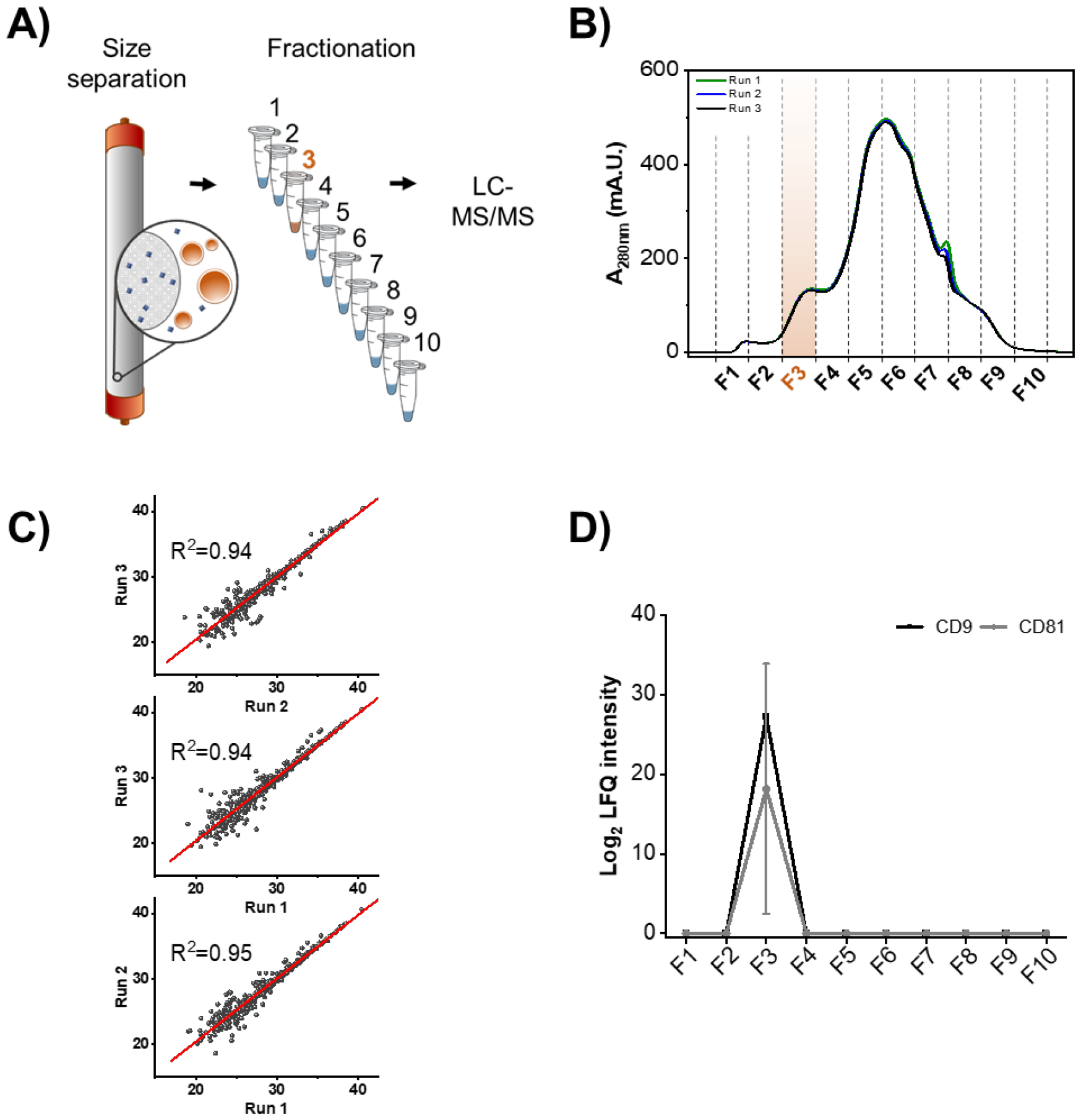
Human plasma extracellular vesicle (EV) fractionation by size exclusion chromatography. (A) Scheme of human plasma fractionation by size exclusion chromatography in a Superose 6 GE 5/150 column. (B) Chromatograms resulting from three consecutive runs. Fraction collection is delimited by dashed lines and the fraction containing EV is highlighted in orange. (C) Scatter plots of Log_2_ intensity of identified proteins comparing the three different runs. Linear regression curves (red) and coefficient of determination (R^2^) are also shown for each plot. (D) Elution profiles of the EV markers cluster of differentiation (CD)81 and CD9 is shown as average and standard deviation from three consecutive runs. Abbreviations: LC-MS/MS – liquid chromatography-tandem mass spectrometry, LFQ – label-free quantification, mA.U. – absorbance units (x10^−3^).

We further tested our pipeline by examining the EV response in 50 µL of plasma from 9 firefighters immediately before and after an acute (45-minute) bout of exercise obtained as described before [8]. Acute exercise has been previously shown to mobilize EV, largely from the skeletal muscle, into circulation [9]; and their protein cargo is known to play an important role in exercise physiology [10]. In the chromatograms, we observed a net increase in the absorbance of fractions corresponding to EV elution in 6 out of 9 individuals post-exercise (**Figure 2**), supporting previous reports of EV mobilization following an acute bout of strenuous exercise [1]. The EV-enriched fractions then underwent acetone precipitation followed by S-trap digestion to further minimize sample loss [11], and were analyzed by data independent acquisition (DIA) on a Q-Exactive HFX mass spectrometer (Thermo Scientific). We build a peptide library by using pooled aliquots of each sample that were submitted to micro-fractionation and data-dependent acquisition (DDA) [12]. Individual samples were analyzed by DIA with tandem mass spectra were collected from the range of 400 to 900 m/z with 10 m/z increment windows (10.0 m/z isolation width; 30% normalized collision energy; 70,000 resolution at 400 m/z). Both DDA and DIA data were analyzed together in FragPipe with similar parameters described above, and quantitative information was extracted with DIA-NN [13]. A total of 838 proteins were identified, of which 261 proteins (32%) were previously identified as EV proteins based off of our meta-analysis (**Supplementary table 3**) [6]. To test the variability of our platform, we randomized, prepared, and analyzed in parallel a commercial human plasma sample, used as a measure of quality control. The quality control sample had a correlation average of 0.91 (**Figure 3A**), indicating low variability of the platform.

**Figure 2.**
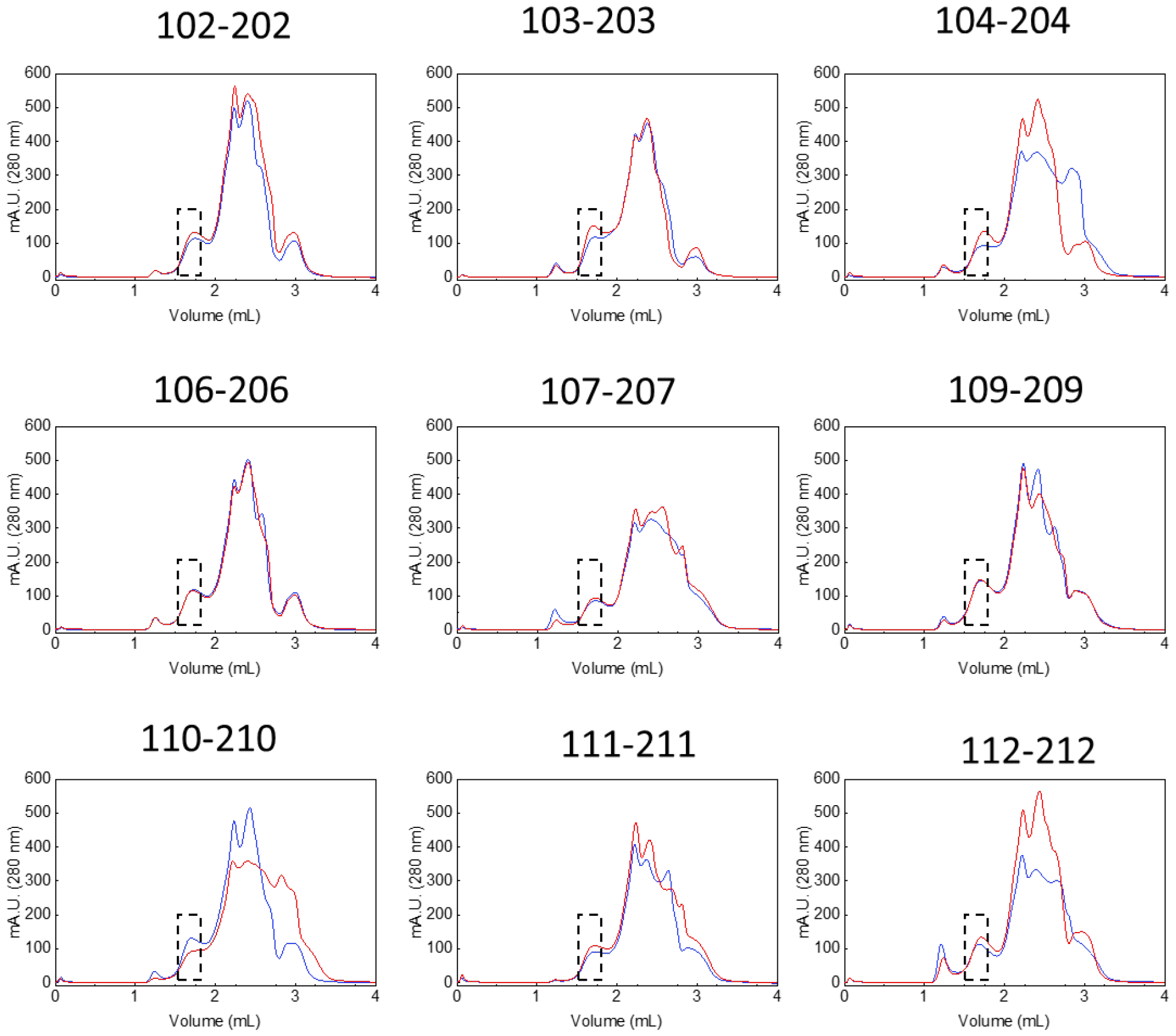
Chromatographic profile of plasma from firefighter volunteers fractionated by size exclusion chromatography. 50 µL of human plasma pre-(BLUE) and post-(RED) exercise were injected into a Superose 6 increase 5/150 and separated in an isocratic gradient at a flow rate of 0.3 mL/min. Fraction number 3 (dashed box) of pre- and post-exercise of each individual was submitted to proteomics analysis.

**Figure 3.**
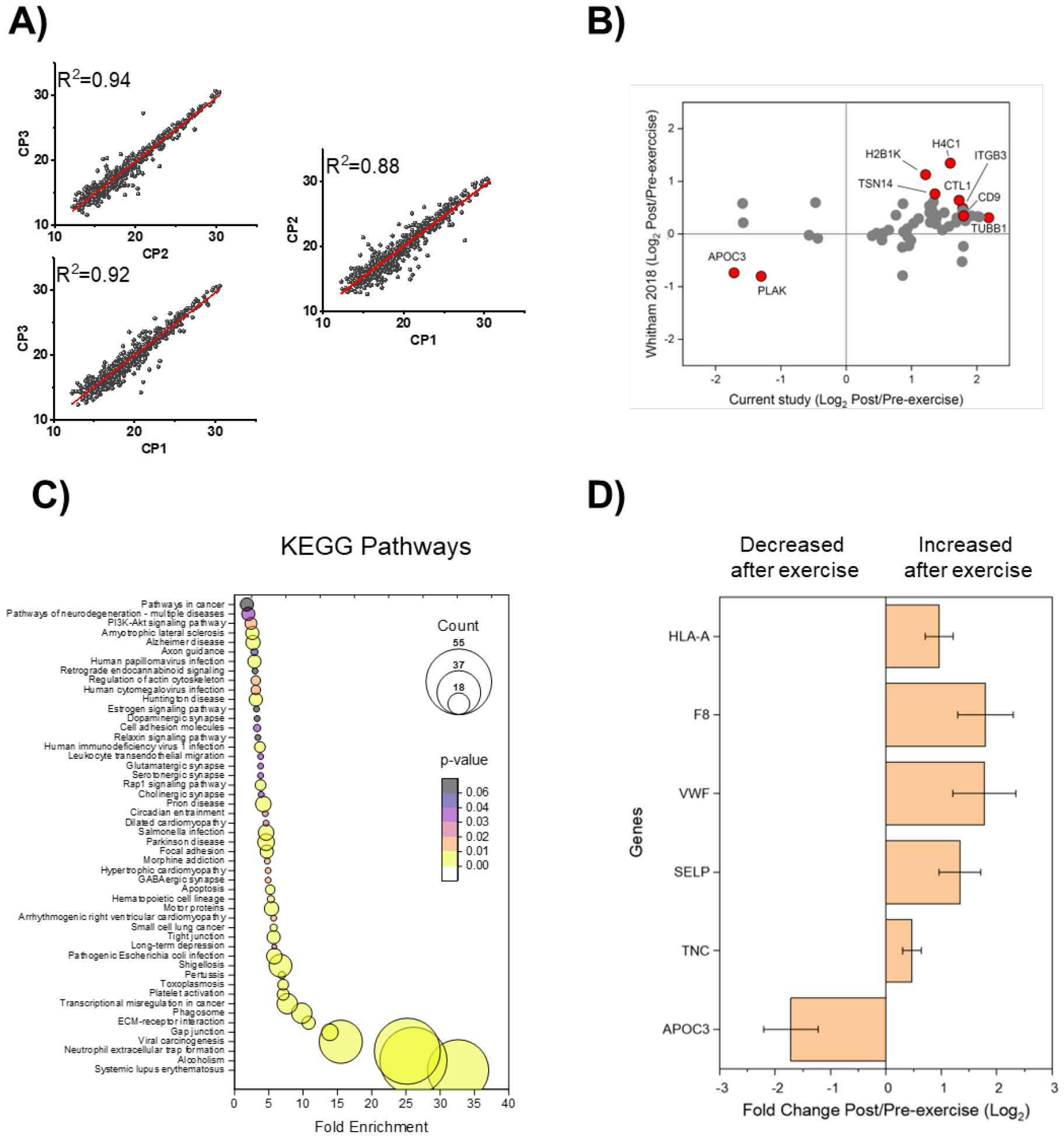
Analysis of plasma extracellular vesicles from a cohort of firefighters that underwent an intense exercise bout. (A) Scatter plot of averaged Log_2_ intensity of identified proteins comparing three commercial plasma (CP) runs, analyzed at beginning, middle and end of the experiment, linear regression curves (red) and coefficient of determination (R^2^) are also shown for each plot. (B) Cross-validation of differentially abundant proteins post-exercise found in common with Whitham *et al*. 2018 [1]. Examples of proteins with similar regulation between both studies are highlighted in red. (C) Functional-enrichment analysis of proteins regulated post-exercise. (D) Examples of regulated proteins that are relevant to exercise physiology: immunoregulatory HLA class I histocompatibility antigen, A (HLA-A); coagulation factor VIII (F8); von Willebran factor (VWF); P-selectin (SELP); tenascin C (TNC) and apolipoprotein C-III (APOC3).

Given the repeatability of our platform, we next examined the utility of our platform to provide biological insight into the acute exercise response via statistical and functional-enrichment analyses. In response to acute exercise, we observed 10 downregulated and 55 upregulated proteins in EV-fractions (p<0.05) (**Supplementary table 4**). We next cross-validated biomarker candidates by re-analyzing the plasma EV proteomic response to acute aerobic exercise from a separate cohort of subjects [1]. Out of the regulated proteins, 57 EV proteins were also detected in the data from the previous work, and 43 proteins displayed a similar temporal response to acute exercise despite differences in exercise regimen. Here we observed increased abundance of the EV marker CD9 and decreased abundance of apolipoprotein C-III (APOC3) (**Figure 3B**). APOC3 is a lipoprotein lipase inhibitor [14] that its downregulation thus permits lipoprotein lipase-mediated fatty acid release to meet the energetic demands of the exercise.

Subsequent functional-enrichment analysis [15] of the 65 exercise-responsive EV proteins in our data set revelated enrichment of 50 pathways following acute exercise (**Figure 3C**), highlighting candidate biological functions for exercise-related EVs. Upregulated pathways included pathways involved in acute inflammatory responses, such as neutrophil activation and coagulation. The acute inflammatory and pro-thrombotic response to strenuous exercise is well documented [16-18]. In agreement, we detected increased abundance of coagulation EV proteins, including coagulation factor VIII (F8), P-selectin (SELP) and von Willebran factor (VWF), immunoregulatory HLA class I histocompatibility antigen, A (HLA-A), and tenascin C (TNC), a marker of muscle damage [19], following acute exercise (**Figure 3D**). These results indicate that our platform can also provide insights on the biological roles of EV.

In this work, we developed a SEC platform for EV fractionation that is fast (15 minutes), robust, has low variability, and dramatically reduces the required sample volume to 50 µL. The application of this platform to a cohort of firefighters that underwent strenuous physical training showed key proteins mobilized in EV in response to acute exercise. We were also able to cross validate biomarker candidates with data from a previous study, demonstrating the potential applicability of this method for clinical studies. We confirmed the method robustness by analyzing samples from multiple individuals. Although this platform has not yet been tested for large scale applications, as the method is currently designed, it could be fully automated, allowing for easy application of the method to large scale studies.

## Abbreviations

(CD9): cluster of differentiation 9
(CD81): cluster of differentiation 81
(DDA): data-dependent acquisition
(DIA): data-independent acquisition
(EV): extracellular vesicles
(FPLC): fast protein liquid chromatography
(SEC): size-exclusion chromatography

## ACKNOWLEDGMENTS

Part of the work was performed in the Environmental Molecular Sciences Laboratory, a U.S. Department of Energy (DOE) national scientific user facility at Pacific Northwest National Laboratory (PNNL) in Richland, WA. Battelle operates PNNL for the DOE under contract DE-AC05-76RLO01830.

## SUPPORTING INFORMATION

**Supplementary Table 1** - Identification and quantification of plasma proteins fractionated by size-exclusion chromatography.

**Supplementary Table 2** – Enrichment of each size exclusion chromatography fraction with previous characterized extracellular vesicles proteins. P-values were calculated using Fisher’s exact test.

**Supplementary Table 3** – Analysis of extracellular vesicles from plasma of a cohort of wildland firefighters that underwent a 45 min intensive physical activity.

**Supplementary Table 4** – Differentially abundant proteins found in the plasma extracellular vesicle fraction from the wildland firefighters that underwent a 45 min intensive physical activity. P-values were calculated using Student’s *t*-test.

## DATA AVAILABILITY

Mass spectrometry data are publicly available in MassIVE repository, a member of the ProteomeXchange Consortium, under the accession number MSV000093427.

## References

[1] Whitham, M., Parker, B. L., Friedrichsen, M., Hingst, J. R., et al., Extracellular Vesicles Provide a Means for Tissue Crosstalk during Exercise. Cell Metab 2018, 27, 237–+.

[2] Liangsupree, T., Multia, E., Riekkola, M. L., Modern isolation and separation techniques for extracellular vesicles. J Chromatogr A 2021, 1636.

[3] Colombo, M., Raposo, G., Théry, C., Biogenesis, Secretion, and Intercellular Interactions of Exosomes and Other Extracellular Vesicles. Annu Rev Cell Dev Bi 2014, 30, 255–289.

[4] Doyle, L. M., Wang, M. Z., Overview of Extracellular Vesicles, Their Origin, Composition, Purpose, and Methods for Exosome Isolation and Analysis. Cells-Basel 2019, 8.

[5] Vergauwen, G., Tulkens, J., Pinheiro, C., Cobos, F. A., et al., Robust sequential biophysical fractionation of blood plasma to study variations in the biomolecular landscape of systemically circulating extracellular vesicles across clinical conditions. J Extracell Vesicles 2021, 10.

[6] Vallejo, M. C., Sarkar, S., Elliott, E. C., Henry, H. R., et al., A proteomic meta-analysis refinement of plasma extracellular vesicles. Scientific Data 2023, 10, 837.

[7] Kong, A. T., Leprevost, F. V., Avtonomov, D. M., Mellacheruvu, D., Nesvizhskii, A. I., MSFragger: ultrafast and comprehensive peptide identification in mass spectrometrybased proteomics. Nat Methods 2017, 14, 513–520.

[8] Nakayasu, E. S., Gritsenko, M. A., Kim, Y. M., Kyle, J. E., et al., Elucidating regulatory processes of intense physical activity by multi-omics analysis. Military Med Res 2023, 10.

[9] Rome, S., Forterre, A., Mizgier, M. L., Bouzakri, K., Skeletal Muscle-Released Extracellular Vesicles: State of the Art. Front Physiol 2019, 10.

[10] Fruhbeis, C., Helmig, S., Tug, S., Simon, P., Kramer-Albers, E. M., Physical exercise induces rapid release of small extracellular vesicles into the circulation. J Extracell Vesicles 2015, 4, 28239.

[11] Zougman, A., Selby, P. J., Banks, R. E., Suspension trapping (STrap) sample preparation method for bottom-up proteomics analysis. Proteomics 2014, 14, 1006–1010.

[12] Dou, M., Tsai, C. F., Piehowski, P. D., Wang, Y., et al., Automated Nanoflow Two-Dimensional Reversed-Phase Liquid Chromatography System Enables In-Depth Proteome and Phosphoproteome Profiling of Nanoscale Samples. Anal Chem 2019, 91, 9707–9715.

[13] Demichev, V., Messner, C. B., Vernardis, S. I., Lilley, K. S., Ralser, M., DIA-NN: neural networks and interference correction enable deep proteome coverage in high throughput. Nat Methods 2020, 17, 41–44.

[14] Ginsberg, H. N., Goldberg, I. J., Broadening the Scope of Dyslipidemia Therapy by Targeting APOC3 (Apolipoprotein C3) and ANGPTL3 (Angiopoietin-Like Protein 3). Arterioscl Throm Vas 2023, 43, 388–398.

[15] Huang da, W., Sherman, B. T., Lempicki, R. A., Systematic and integrative analysis of large gene lists using DAVID bioinformatics resources. Nat Protoc 2009, 4, 44–57.

[16] Quindry, J. C., Stone, W. L., King, J., Broeder, C. E., The effects of acute exercise on neutrophils and plasma oxidative stress. Med Sci Sports Exerc 2003, 35, 1139–1145.

[17] Nieman, D. C., Pence, B. D., Exercise immunology: Future directions. J Sport Health Sci 2020, 9, 432–445.

[18] Womack, C. J., Nagelkirk, P. R., Coughlin, A. M., Exercise-induced changes in coagulation and fibrinolysis in healthy populations and patients with cardiovascular disease. Sports Med 2003, 33, 795–807.

[19] Ekstrom, O., Strom, K., Mir, B. A., Laurila, E., et al., Increasing circulating levels of Tenascin C in response to the Wingate anaerobic test. Clin Physiol Funct Imaging 2023, 43, 271–277.

